# Regularity extraction under stress: boosted statistical learning but unaffected sequence learning

**DOI:** 10.1101/2020.05.13.092726

**Authors:** Eszter Tóth-Fáber, Karolina Janacsek, Ágnes Szőllősi, Szabolcs Kéri, Dezső Németh

## Abstract

Acute stress can crucially influence learning and memory processes. One of the key processes underlying human learning and memory is the ability of our brain to rapidly detect and extract regularities from sensory input across time and space leading to effective predictive processing. Here, we aimed to get an in-depth look into the effect of stress on the acquisition of two aspects of regularity extraction. We examined whether and how stress affects (1) processes and learning of probability-based, statistical regularities and (2) serial order-based, sequential regularities in the same experimental design, and (3) explicit access to the acquired information. Considering that the acquisition of statistical regularities is a rapid process, we primarily focused on the early phases of the task. We induced stress with the Socially Evaluated Cold Pressor Test in 27 young adults, while 26 participants were enrolled in the control group. Salivary cortisol levels and subjective ratings of affective states showed successful stress induction. After the stress induction, we measured regularity extraction with the cued Alternating Serial Reaction Time task. We found that stress promoted the acquisition of statistical regularities and did not alter the learning of sequential regularities. Post-block reports showed weaker explicit access to the sequential regularities in the stress group. Our results give a process-level understanding on how stress alters learning and memory functions related to predictive processes.

## 1. Introduction

Stressful situations are ubiquitous parts of our everyday life. Acute stress leads to changes in both cognitive and affective processes (Joëls et al., 2011, Schwabe et al., 2012; Ehlers and Todd, 2017, Dolfen et al., 2019). Previous studies have shown the crucial influence of stress on learning and memory functions (e.g., Kirschbaum et al., 1996; Nater et al., 2007; Schwabe & Wolf, 2012). One of the key processes underlying human learning and memory is the ability of our brain to rapidly extract regularities from sensory input across time and space (Perruchet & Pacton, 2006, Frost et al., 2015, Saffran & Kirkham, 2018, Conway, 2020). This knowledge is then used to update our predictions of future events (Misyak et al., 2010, Fine & Jaeger, 2013). The aim of the present study is to determine how stress affects two aspects of such regularity extraction.

The available information in our environment is diverse, we can detect and extract several different kinds of regularities, from simple structures such as deterministic rules (i.e., sequential, serial order-based regularities) to complex regularities such as statistical, probability-based regularities (Németh et al., 2013; Kóbor et al., 2018; Simor et al., 2019). The latter one termed statistical learning, referring to the acquisition of shorter-range relations between visual stimuli that is mainly based on statistical regularities (i.e., differentiating between more and less probable stimuli). The extraction of sequential regularities termed sequence learning, referring to the acquisition of a series of stimuli that repeatedly occur in the same deterministic order (often intermixed with random stimuli, resulting in an alternating sequence structure). Importantly, from a theoretical perspective, sequence learning can be viewed as a form of statistical learning. Considering transitional probabilities, statistical learning refers to the learning of second-order transitional probabilities that are less than one, while sequence learning refers to the learning of second order transitional probabilities that are equal to one. Although both can be considered as processes to acquire transitional probabilities, statistical and sequence learning are distinguishable on both neural and behavioral levels: they develop differently at the level of event-related potentials (Kóbor et al., 2018), and they show distinct neural oscillations during consolidation (Simor et al., 2019). Crucially, on the behavioral level, participants acquire statistical regularities rapidly and show consistent, stable performance thereafter (Kóbor et al., 2018; Simor et al., 2019). Simor et al. (2019) explored the trajectory of statistical learning in detail. The results showed that picking up statistical regularities from the stimulus stream happens very quickly and it is not due to pre-existing, not learning-related tendencies. The results also indicated that additional training helps strengthening the acquired statistical regularities. In contrast to statistical learning, acquiring sequential regularities requires more time, suggested by the gradual trajectory of sequence learning (Kóbor et al., 2018; Simor et al., 2019).

To the best of our knowledge, only a handful of studies investigated the effect of stress on the acquisition of statistical or sequential regularities (Römer et al., 2011; King, 2013; Dolfen et al., 2019). In King’s (2013) study, participants were exposed to a statistical regularity without being aware that learning were taking place. Stressed participants showed overall enhanced learning in accuracy. Regarding the acquisition of sequential regularities, Römer et al. (2011) examined the effect of exogenous cortisol administration on sequence learning. While both the cortisol and placebo groups showed significant sequence learning, cortisol changed the time course of learning. The cortisol group showed delayed learning with decreased performance at the beginning of the task and equaled performance at the end of the task. In contrast, Dolfen et al. (2019) did not find any effect of experimental stress exposure on deterministic motor sequence learning using a bimanual finger-tapping task. To sum up, only three studies examined whether stress influences the extraction of statistical or sequential regularities.

Altogether, the abovementioned prior studies do not allow us to draw strong conclusion about whether and how stress influences the acquisition of different types of regularities that are extracted from the environment. The aim of the present study thus was to test the effect of stress on two aspects of regularity extraction, namely the acquisition of probabilistic and order-based information (i.e., statistical learning in a narrow sense and sequence learning). As prior studies are scarce and inconclusive, we use an exploratory approach to investigate this question. As statistical learning is a very rapid process (Frost et al., 2015; Kóbor et al., 2018; Simor et al., 2019; Conway, 2020), if stress affects this aspect of learning, we expect it to occur at the beginning of the task, irrespective of whether stress diminishes or enhances the acquisition. In the case of sequence learning, as it typically shows a gradual trajectory (Kóbor et al., 2018; Simor et al., 2019), if stress affects this aspect of learning, we expect the alterations to occur in the later stages of the task.

## 2. Materials and methods

### 2.1. Participants

Sixty-five healthy undergraduates enrolled in our study. Two participants’ baseline salivary cortisol levels were more than three standard deviations away from the mean of the sample. Therefore, these two participants were considered as outliers, and their data were not included in the analyses. Three participants were also excluded from the analyses because they did not provide sufficient saliva for cortisol analysis. Additionally, as following instructions in the cued ASRT task (for details, see section 2.4) is crucial, seven participants were excluded as they did not meet this requirement (for details, see section 2.5; for analysis on the sample containing these participants, see Supplementary Material).

The final sample consisted of 53 participants (15 men, 38 women; M_age_ = 20.9 years, SD = 1.7). Participants were randomly assigned to either the control (7 men, 19 women) or the stress (8 men, 19 women) group. 10 women in the stress group and 7 women in the control group took oral contraceptives regularly. All participants had normal or corrected-to-normal vision, none of them reported a history of any psychiatric, neurological or any other chronic medical problems. They performed in the normal range of standard neuropsychological tests (Counting Span task: M_stress_ = 3.87, M_control_ = 3.73; Digit Span task: M_stress_ = 7.03, M_control_ = 7.23), with no difference between the stress and control groups (all *<u>p</u>*s > .537, Cohen’s *d* < 0.176).

Participants were recruited from a university course in exchange for course credit. All participants provided written informed consent before enrolment. The study was approved by the research ethics committee of the Eötvös Loránd University in Hungary and was conducted in accordance with the Declaration of Helsinki.

### 2.2. Experimental design

To exclude the effect of critical factors on salivary cortisol levels, participants were asked not to eat, drink (except for water), smoke, and make physical exercise two hours prior to the experiment (see Kirschbaum and Hellhammer, 1994; Lovallo et al., 2006; Badrick, Bobak, et al., 2007; Badrick et al., 2007). To avoid interference with the cortisol circadian cycle (see Clow et al., 2010), assessments took place between 1 and 7 p.m.

Each experimental session started with a 15-min preparatory phase while participants remained in a preparatory room for 15 minutes to minimize the effect of potentially stress-inducing factors (such as new environment; see Lupien et al., 2007). Then, participants completed a practice session (3 blocks, ca. 3 minutes) of the regularity extraction task. Note that in the practice session stimulus appeared randomly, hence it could not influence later statistical and sequence learning. To measure cortisol levels, saliva samples were collected at different time points during the experimental session (see Figure 1). The first saliva sampling was conducted immediately after the practice session of the regularity extraction task. Baseline saliva sampling was followed by the stress induction or control task (see the details below). To assess subjective stress levels, immediately after the stress induction or the control task, participants were required to rate how stressful (0 = not at all, 100 = very stressful), painful (0 = not at all, 100 = very painful), and unpleasant (0 = not at all, 100 = very unpleasant) the stress inducing or the control task was (Table 1). The second saliva sample was collected 15 minutes after stressor offset/the end of the control task, followed by the regularity extraction task. The third saliva sample was collected immediately after the regularity extraction task (approximately 40 minutes after stressor offset/the end of the control task). Samples were collected using Eppendorf Safe-Lock Tubes (1.5 mL), were kept at −20 °C between the experimental session and the analysis. Saliva samples were analyzed using Salimetrics immunoassays.

**Table 1.** Descriptive data of the participants’ subjective stress measurements.

|  | Group |  |  |  | *t* | *p* |
| --- | --- | --- | --- | --- | --- | --- |
|  | Stress |  | Control |  |  |  |
|  | (*n* = 27) |  | (*n* = 26) |  |  |  |
|  | *M* | *SD* | *M* | *SD* |  |  |
| Subjective ratings of stress | 29.88 | 26.14 | 8.34 | 12.43 | -3.85 | <.001** |
| Subjective ratings of unpleasantness | 56.29 | 28.78 | 11.30 | 14.29 | -7.24 | <.001** |
| Subjective ratings of painfulness | 58.14 | 25.56 | 1.34 | 5.92 | -11.23 | <.001** |

*Note.* Immediately after the stress induction or control task, participants rated how stressful, unpleasant, and painful (where 0 = not at all, 100 = very) the stress induction or control task was. \*\* $p < .001$ .

**Figure 1.**
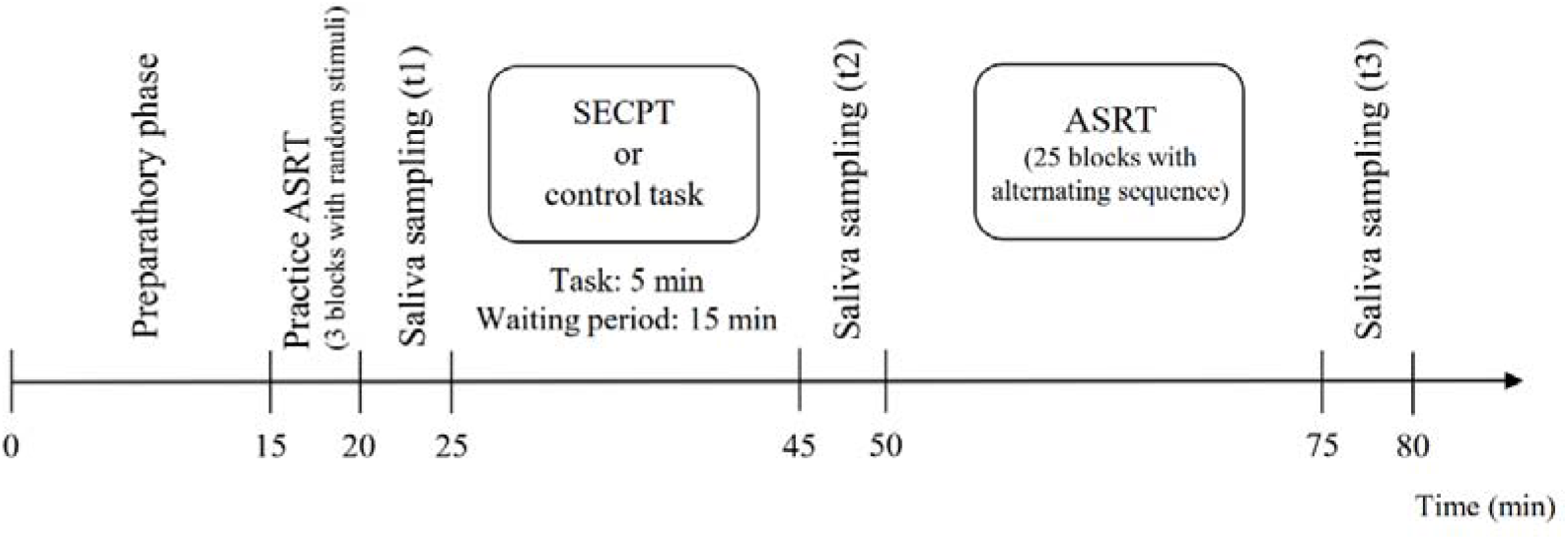
Experimental procedure. Cortisol levels were assessed from saliva at three time points: immediately before the stress induction or control task (t1), 15 minutes after the stress induction or control task (t2), and immediately after the ASRT task, ca. 45 minutes after the stress induction or control task (t3). ASRT: Alternating Serial Reaction Time task; SECPT: Socially Evaluated Cold Pressor Test.

### 2.3. Stress induction task and control condition

Individuals in the stress group participated in the Socially Evaluated Cold Pressor Test (Schwabe et al., 2008). This task was developed to induce moderate stress in laboratory settings using both physical and psychosocial stressors. Participants were instructed to put their hand into cold water (0-3 °C) for three minutes while a female experimenter observed their behavior. The observer did not interact with the participants, just passively monitored their behavior. Participants were also told that video recording would be made to later analyze their non-verbal behavior. They were informed that they could interrupt the task if it would be too painful or too uncomfortable.

There were no stress inducing factors (i.e. cold water, observer, and video recorder) in the control condition. Control participants were required to put their hand into warm water (35-37 °C) for three minutes. A female experimenter stayed in the experimental room (just as in the stress condition), however, this time she did not observe participants’ behavior.

### 2.4. Regularity extraction task

Regularity extraction was measured by the cued version of the Alternating Serial Reaction Time (ASRT) task (Song et al., 2007a; Németh et al., 2013; Simor et al., 2019). In this task, a target stimulus (either a dog’s head or a penguin) appeared in one of the four possible locations, which were horizontally arranged and equally spaced empty circles. Participants were asked to press the corresponding key on the keyboard (Z, C, B, or M) as accurately and as fast as they could, using their index-and middle-fingers of both hands. The stimulus remained on the screen until the participant’s response, then, after a 120 ms delay, the next target appeared (note that in this task, one stimulus corresponds to one trial).

Participants started the experimental session with a practice session of the ASRT task consisting of 3 blocks with 85 trials each. Importantly, in the practice session, stimulus appeared randomly, *without* any predetermined sequence. In the task, however, the presentation of the stimuli followed an 8-element alternating sequence where pattern and random trials alternated with each other (e.g. 1-r-2-r-4-r-3-r, where numbers represent the predetermined locations from left to right and ‘r’ indicates a randomly selected location). The pattern and random trials were marked differently, pattern trials indicated by the dog’s head, and random trials represented by the penguins. Participants were told that the appearance of the dogs always followed a predetermined sequence, while penguins always appeared on a randomly chosen location. They were instructed to find the pattern defined by the dog’s appearance to improve their performance. They had no information about the length of the predetermined sequence. The alternation of pattern and random trials creates six distinguishable sequence permutations: 1-r-2-r-3-r-4-r, 1-r-2-r-4-r-3-r, 1-r-3-r-2-r-4-r, 1-r-3-r-4-r-2-r, 1-r-4-r-2-r-3-r, and 1-r-4-r-3-r-2-r. One of these permutations was selected for each participant and it was counterbalanced between groups and also across participants in each group.

The task contained 25 blocks and each block consisted of 85 trials. The first five trials of each block were random trials for practice, then the unique 8-element predetermined sequence was presented 10 times. After each block, participants were instructed to type the order of the dog’s head using the corresponding keys to measure awareness of the sequence structure. The *post-block sequence report* lasted until the participants gave 12 consecutive responses, which are ideally the 4-element sequence three times. For these post-block sequence reports, we calculated how many responses out of the total 12 were correct after each block, expressed in terms of percentages. The mean of these block-level values was calculated for each participant and used as a measure of their *explicit sequence knowledge* acquired during the task.

The alternating sequence in the ASRT task makes some runs of three successive trials – referred to as *triplets* – more probable than others. For example, if the sequence is 1-r-2-r-4-r-3-r, triplets such as 1-X-2, 2-X-4, 4-X-3, 3-X-1 (where X indicates the middle element of the triplet) occur with a higher probability since their last trial can be either predetermined or random. However, 3-X-2 or 4-X-2 occur with a lower probability as the third trial could only be random. The more probable triplet types are hereinafter referred to as high-probability triplets and the latter types as low-probability triplets (Howard and Howard, 1997; Németh et al., 2013). Since high-probability triplets can end with a pattern trial due to the predetermined sequence or with a random trial by chance, we can differentiate between pattern high-probability and random high-probability triplets. Low-probability triplets can only end with a random trial, as pattern trials are always at high probability. Note that each element (i.e., trial) was categorized as the third element of either a high-probability or a low-probability triplet. In sum, three trial types can be differentiated: (1) trials that belong to the predetermined sequence and are the final element of a high-probability triplet labeled as *pattern trials*, (2) trials that occur randomly and also are the final element of a high-probability triplet called *random high trials*, and (3) random elements that appear as the last element of a low-probability triplet called *random low trials*.

Based on these trial types, we can differentiate between two types of regularity extraction: statistical learning and sequence learning (Figure 2C; Howard & Howard, 1997; Németh et al., 2013; Kóbor et al., 2018; Simor et al., 2019). *Statistical learning* refers to the acquisition of probability-based, statistical regularities, and it is measured by the difference in reaction times (RTs) between random high and random low trials. These trials share the same sequence structure as they are both random trials but differ in statistical properties as they correspond either to the final trial of a high-probability or a low-probability triplet. Greater statistical learning is indicated by faster RTs on random high than on random low trials.

**Figure 2.**
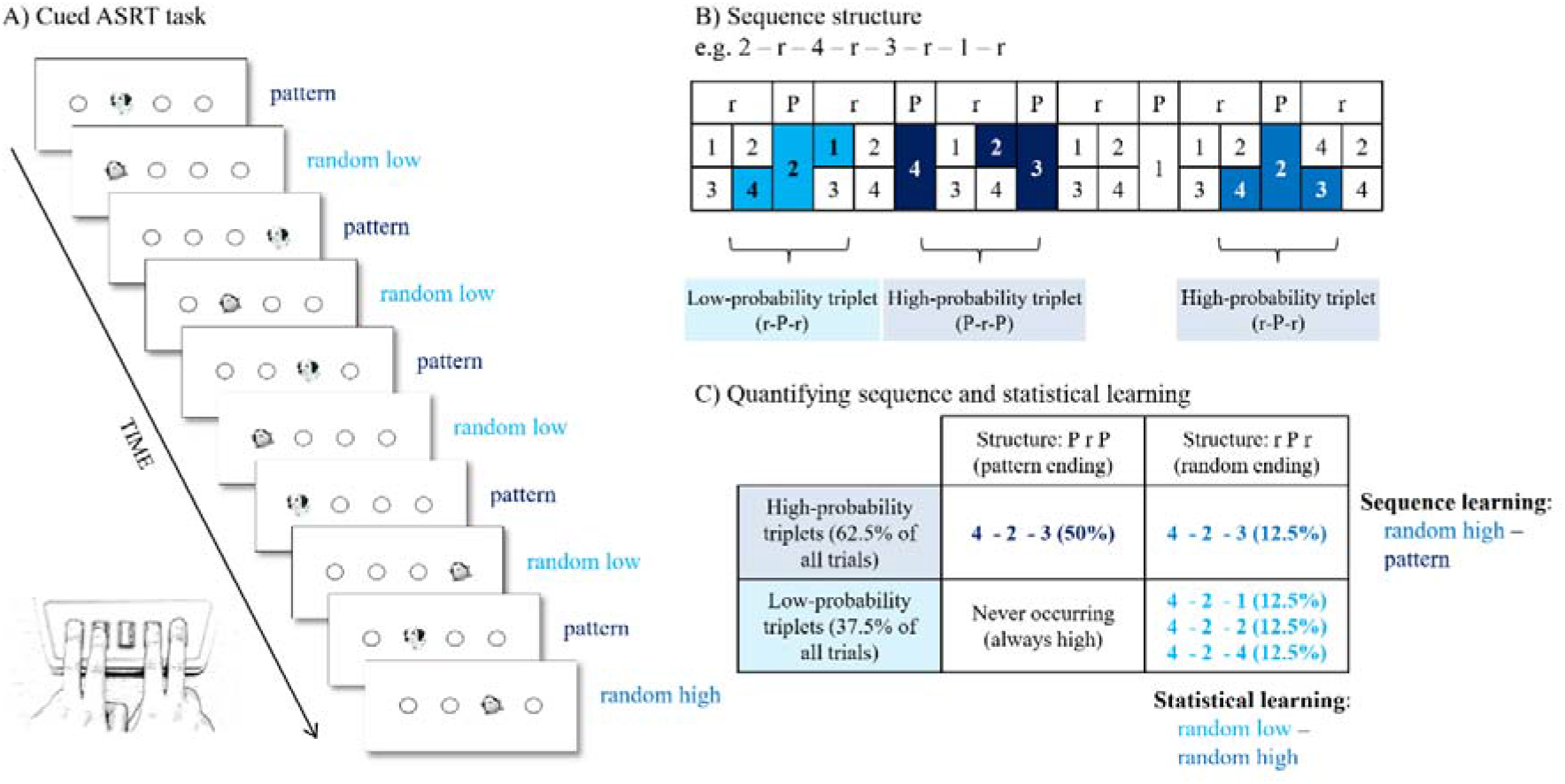
The cued Alternating Serial Reaction Time (ASRT) task. (A) Pattern and random trials were presented in an alternating fashion in the ASRT task. Pattern trials were indicated by a picture of a dog and random trials were indicated by a picture of a penguin. (B) An example of the alternating sequence structure. Here, numbers indicate pattern trials and ‘r’ indicates a randomly selected location. The alternating sequence makes some runs of three consecutive trials (called triplets) more probable than others, termed high- and low-probability triplets, respectively. Among high-probability triplets, we determined pattern high-probability triplets (dark blue shading in panel B and dark blue font in panel C) and random high-probability triplets (blue shading in panel B and blue font in panel C). Low-probability triplets can occur only in random positions (light blue shading in panel B and light blue font in panel C). (C) The underlying learning processes in the task. Sequence learning is quantified by contrasting the RTs on the pattern and random high trials. Statistical learning is quantified by contrasting the RTs on the random high and random low trials. Adapted from Németh et al. (2013).

*Sequence learning* refers to the acquisition of serial order-based, sequential regularities, and it is measured by the difference in RTs between pattern and random high trials. These trials share the same statistical properties as they are both the last element of a high-probability triplet, but differ in sequence properties as pattern trials are part of the predetermined sequence, while random high trials appear on a random location. Greater sequence learning is defined as faster RTs on pattern than on random high trials.

### 2.5. Statistical analysis

Statistical analyses were carried out by SPSS version 25.0 software (SPSS, IBM). As participants were instructed to find the alternating sequence and report that during the task, performance on the sequence report (calculated as the average percentage of correctly reporting the sequence after the ASRT task blocks) could show whether participants adequately followed the instructions of the task. In the present study, seven participants exhibited below 50% performance, suggesting that they did not follow the instructions properly and were therefore excluded from the analyses (see also section 2.1; for analysis on the sample containing these participants, see Supplementary Material).

Statistical analysis of the effectiveness of stress induction was the same as in prior studies (e.g. Guenzel et al., 2014, Szőllősi et al., 2017). We conducted a mixed design ANOVA on cortisol levels as dependent variable, with TIME (t1, t2, and t3) as a within-subject factor, where t1 = immediately before stress/control procedure, t2 = 15 minutes after stress/control procedure, t3 = 45 minutes after stress/control procedure, and with STRESS EXPOSURE (stress or control) as a between-subject factor. The effectiveness of stress induction was also quantified with subjective variables. Immediately after the stress induction or control task, participants were asked to rate how stressful, painful, and unpleasant the stress induction or the control task was. We compared these subjective ratings of affective state between the stress and control groups with independent sample t-tests.

To assess participants’ average speed prior to stress induction or control task, and to exclude the possibility of pre-existing group differences influencing the effect of stress on learning, we conducted a mixed design ANOVA on the practice session of the ASRT task. This session consisted of three blocks and stimuli presented randomly, therefore average RTs could be assessed. The mixed design ANOVA involved GROUP (stress vs control) as a between-subject factor and BLOCK (1-3) as a within-subject factor.

Statistical analysis of the ASRT task was based on previous studies (Németh et al., 2013; Kóbor et al., 2018; Simor et al., 2019). Two types of low-probability triplets, namely repetitions (e.g. 111, 222) and trills (e.g. 121, 242) were excluded from the analysis, since participants often show pre-existing tendencies to these trials (Song et al., 2007a, 2007b). Following prior studies (e.g. Németh et al., 2013; Janacsek et al., 2018; Kóbor et al., 2017; 2018; Simor et al., 2019; Takács et al., 2018), the ca. 1-minute-long blocks were organized into larger segments labeled as epochs. Each epoch consists of five blocks, thus, we analyzed five epochs of the ASRT task.

As described in section 2.4, each trial was determined as the last element of a *pattern high, random high*, or *random low* triplet. The median of RT data for correct responses was calculated for each participant in each epoch, separately for the three trial types. As presented above, two aspects of regularity extraction, sequence learning and statistical learning can be assessed by the ASRT task (Németh et al., 2013; for further details, see Introduction and the task description above). To examine the two aspects, RT data were analyzed in a mixed design ANOVA. At first, as statistical learning is a rapid process (Kóbor et al., 2018; Simor et al., 2019), we focused on the early phase of the task. We used a mixed design ANOVA with PROBABILITY (random high vs random low trials) as a within-subject factor and GROUP (stress vs control) as a between-subject factor on the RT data showed in the first epoch. Similarly, at first, sequence learning was compared only in the last epoch of the task as this learning process shows a gradual trajectory (Kóbor et al., 2018; Simor et al., 2019), and therefore, the effect of stress may emerge in this later phase of learning. Hence, first, we used a mixed design ANOVA with ORDER (pattern vs random high trials) as a within-subject factor and GROUP (stress vs control) as a between-subject factor on the RT data showed in the last epoch.

Next, to investigate learning in the learning phase (i.e., on the whole task), statistical learning was quantified with a mixed design ANOVA with PROBABILITY (random high vs random low trials) and EPOCH (1-5) as within-subject factors and GROUP (stress vs control) as a between-subject factor. Similarly, sequence learning was quantified with a mixed design ANOVA with ORDER (pattern vs random high trials) and EPOCH (1-5) as within-subject factors and GROUP (stress vs control) as a between-subject factor. Pairwise comparisons were performed using LSD (Least Significant Difference) to control for type 1 error. The Greenhouse-Geisser epsilon correction was used when necessary. Original *df* values and corrected *p* values (if applicable) are reported together with partial eta-squared (*η*^*2*^ _*p*_) as a measure of effect size.

The analysis on statistical learning showed that, on average, the stress group showed faster RTs compared to the control group. Therefore, we transformed the raw RT data into standardized RTs (for a detailed description, see Results). In conjunction with the analyses on raw RT data, we also present identical analysis on standardized RT data to ensure that potential group differences are not masked by the group differences in average RTs.

## 3. Results

### 3.1. The effectiveness of stress induction

We used both objective (cortisol levels) and subjective (questionnaire-based rating) measurements to test the effectiveness of stress induction. For the *objective measurement*, the ANOVA confirmed the effectiveness of the stress induction. We found a significant TIME × STRESS EXPOSURE interaction (*F*(2, 102) = 29.75, *p* < .001, η^*2*^ _*P*_ *=* .368). The post hoc analysis revealed that cortisol levels in the stress and control groups differed 15 minutes after the stress/control procedure (t2; *M*_stress_ = 32.62 and *M*_control_ = 22.26, respectively, *p* = .048), while immediately before the stress/control procedure (t1) and immediately after the ASRT task (t3) the two groups’ cortisol levels did not differ significantly (both *p*s > .34, see Figure 3). For the *subjective ratings* of affective state after stress induction or control task, the t-tests showed significantly higher subjective stress, pain, and unpleasantness levels in the stress group compared to the control group (see Table 1).

**Figure 3.**
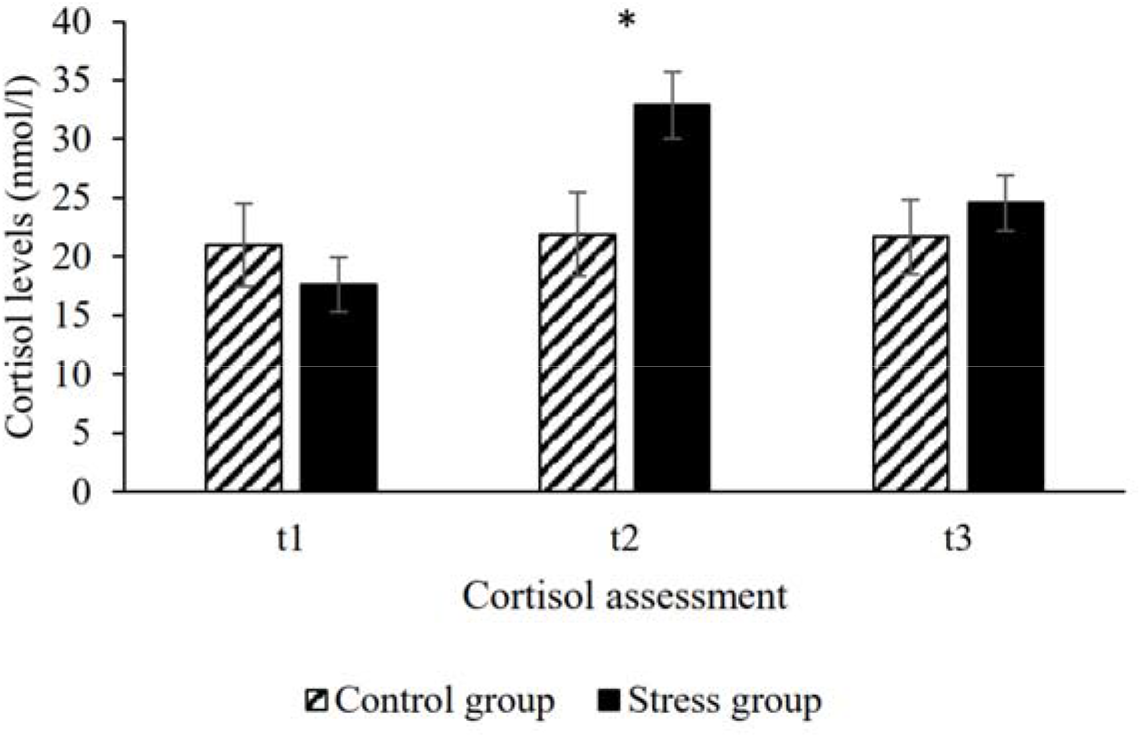
The effectiveness of stress induction. Salivary cortisol levels were assessed immediately before (t1), 15 minutes after (t2), and 45 minutes after (t3) the stress induction or control task. Error bars denote the standard error of the mean (SEM).

### 3.2. Average RTs in the practice session

To compare the two groups’ average speed in the practice session of the ASRT task, a mixed design ANOVA was conducted. The ANOVA revealed that participants became faster as the task progressed (main effect of BLOCK, F(2, 102) = 59.11, p < .001, η^*2*^_*P*_= .54). Importantly, no group differences were found either in average RTs (main effect of GROUP, F(1, 51) = 0.17, p = .68) or in the speed-up during the practice blocks (GROUP × BLOCK interaction, F(2, 102) = 0.29, p = .72).

### 3.3. The effect of stress induction on statistical learning

As statistical learning is a rapid process (Kóbor et al., 2018; Simor et al., 2019), we focused on the beginning of the task. Therefore, we first examined statistical learning only in the first epoch of the learning phase using a mixed-design ANOVA (see section 2.5) between the stress and control groups. Overall, there was no difference in average RTs between the two groups (main effect of GROUP, *F*(1, 51) = 2.67, *p* = .11). Participants responded faster to random high than random low trials (shown by the significant main effect of PROBABILITY, *F*(1, 51) = 32.34, *p* < .001, η^*2*^_*p*_*=* .39), indicating significant statistical learning. Importantly, the stress group showed greater statistical learning than the control group (indicated by the significant PROBABILITY × GROUP interaction, *F*(1, 51) = 4.07, *p* = .05, _η_*2*_*p*_ *=* .07; stress group: *M*_*random high*_ = 373.07 ms, *M*_*random low*_ = 390.59 ms, statistical learning score: *M* = 17.52 ms; control group: *M*_*random high*_ = 401.69 ms, *M*_*random low*_ = 410.04 ms, statistical learning score: *M* = 8.35 ms).

To explore the trajectory of statistical learning in the stress and control groups, we conducted a mixed-design ANOVA on the RTs over the entire learning phase (see section 2.5). This analysis revealed significant differences in average RTs between the two groups with the stress group showing, on average, faster RTs (*M* = 371.19 ms) than the control group (*M* = 391.46 ms) (shown by the significant main effect of GROUP, *F*(1, 51) = 3.98, *p* = .05, _η^*2*^ *p*_ *=* .072). Participants became faster with practice, irrespective of trial type (shown by the significant main effect of EPOCH, *F*(4, 204) = 4.30, *p* = .026, _η_^*2*^_*p*_ *=* .078). Importantly, RTs were faster on random high trials than on random low trials (indicated by the significant main effect of PROBABILITY, *F*(1, 51) = 139.46, *p* < .001, _η_^*2*^_*p*_ *=* .73), revealing significant statistical learning. None of the interactions reached significance (all *p*s > .12, see Table 2).

**Table 2.** Summary of results from ANOVAs performed on raw RT and standard RT data considering statistical learning.

|  | raw RT |  |  | standard RT |  |  |
| --- | --- | --- | --- | --- | --- | --- |
| | $F$ | $p$ | $\eta^2_p$ | $F$ | $p$ | $\eta^2_p$ |
| GROUP | 3.98 | .05* | .07 | < 0.001 | .98 | < .001 |
| PROBABILITY | 139.46 | < .001** | .73 | 142.05 | < .001** | .74 |
| EPOCH | 4.30 | .03* | .08 | 3.31 | .05* | .06 |
| GROUP × PROBABILITY | 1.68 | .20 | .03 | 3.45 | .07+ | .06 |
| GROUP × EPOCH | 0.70 | .46 | .01 | 0.81 | .42 | .02 |
| PROBABILITY × EPOCH | 1.91 | .12 | .04 | 1.70 | .16 | .03 |
| GROUP × PROBABILITY × EPOCH | 1.57 | .19 | .03 | 1.38 | .24 | .03 |

Note. \*\* $p < .01$ , \* $p < .05$ , + $p < .10$ .

The group difference in average RTs revealed by this analysis could affect potential group differences in statistical learning as well, for example, by masking differences in learning as the stress group had less room to improve on the task due to generally faster RTs. To control for the differences in average RTs, we transformed the data in the following way. We divided each participants’ raw RT values of each trial type and each epoch by their own average performance of the first epoch of the task (for a similar approach, see Nitsche et al., 2003; Horváth et al., 2019), then we multiplied these values by 100 for easier presentation and interpretability. Participants’ performance was around 100 at the beginning of the task and changed as the task progressed. On these standardized RT data, we conducted identical analyses as on the raw RT data.

Firstly, we compared learning in the first epoch using a mixed-design ANOVA. The results were identical to the ones on raw RT data. Overall, the groups showed comparable average RTs (main effect of GROUP, F(1, 51) = 1.13, p = .29). Participants responded faster on random high than on random low trials (shown by the main effect of PROBABILITY, F(1, 51) = 32.59, p < .001, η^*2*^_*p*_ *=* .40), indicating significant statistical learning. Moreover, the stress group showed greater learning in the first epoch than the control group (shown by the significant PROBABILITY × GROUP interaction, *F*(1, 51) = 4.46, *p* = .04, _η^*2*^ *p*_ *=* .08).

We also tested the trajectory of statistical learning on the whole task using the standardized RT data. The previously seen group difference in average RTs has been controlled for by the standardization of the data (confirmed by the non-significant main effect of GROUP, *F*(1, 51) < .001, *p* = .98). Identically to the results on raw RT data, analysis showed that participants became faster as the task progressed, irrespective of trial types (main effect of EPOCH, *F*(4, 204) = 3.31, *p* = .05, _η_^*2*^ *p =* .06) and showed significant statistical learning (main effect of PROBABILITY, *F*(1, 51) = 142.05, *p* < .001, _η_^*2*^ *p =* .74). Crucially, this analysis revealed that, overall, the stress group showed marginally greater statistical learning than the control group (as indicated by the marginally significant GROUP × PROBABILITY interaction, *F*(1, 51) = 3.45, *p* = .07, _η_^*2*^_*p*_ *=* .06; stress group: *M*_*random high*_ = 96.34%, *M*_*random low*_ = 101.61%, standardized statistical learning score: *M* = 5.27%; control group: *M*_*random high*_ = 97.04%, *M*_*random low*_ = 100.89%, standardized statistical learning score: *M* = 3.85%). For further details on other interactions, see Table 2.

**Figure 4.**
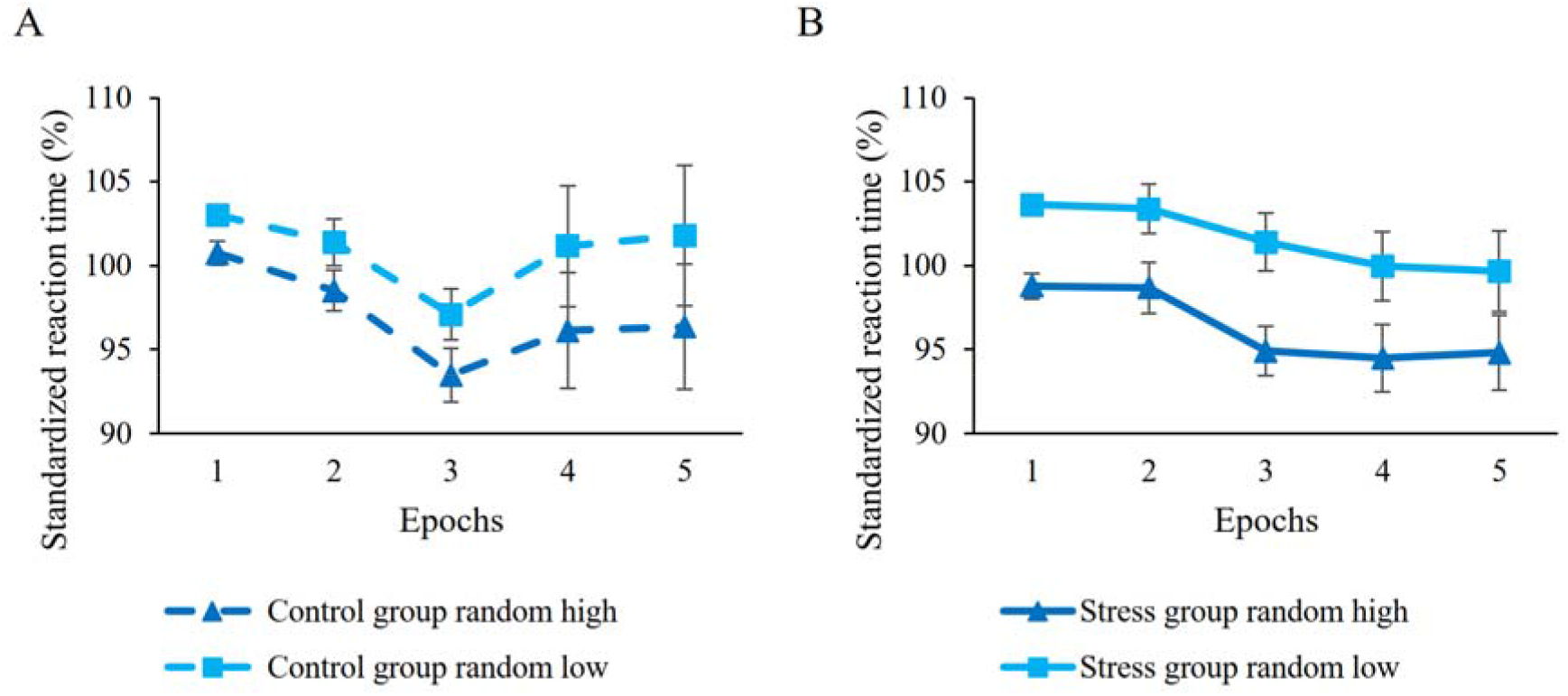
Statistical learning in (A) control and (B) the stress groups. Dashed lines represent the control group, continuous lines represent the stress group. Blue lines with triangle symbols indicate standardized reaction times on the random high trials, light blue lines with square symbols indicate standardized reaction times on the random low trials. Statistical learning is quantified by the distance between dashed and continuous lines, greater distance represents better learning. Error bars denote the SEM.

### 3.4. The effect of stress induction on sequence learning

As sequence learning is a gradual process (Kóbor et al., 2018; Simor et al., 2019), at first, we investigated sequence learning between the groups in the last epoch of the learning phase using a mixed-design ANOVA (see section 2.5). Overall, participants showed similar average reaction times (main effect of GROUP, *F*(1, 51) = 1.21, *p* = .28). Participants responded faster to pattern than to random high trials (main effect of ORDER, *F*(1, 51) = 9.54, *p* = .003, _η^*2*^ *p*_ *=* .16), indicating significant sequence learning. The analysis showed comparable sequence learning between the groups (shown by the non-significant ORDER × GROUP interaction, *F*(1, 51) = 0.76, *p* = .39).

To examine the trajectory of sequence learning on the whole task, we also used mixed-design ANOVA on the RT (see section 2.5). Average RTs did not differ between the groups (main effect of GROUP, *F*(1, 51) = 2.68, *p* = .11). With practice, participants showed faster RTs on both trials (shown by the significant main effect of EPOCH, *F*(4, 204) = 35.35, *p* < .001, _η^*2*^ *p*_ *=* .409). Moreover, participants showed faster RTs on pattern trials compared to random high trials (shown by the main effect of ORDER, *F*(1, 51) = 14.48, *p* < .001, _η_^*2*^_*p*_ *=* .221), indicating significant sequence learning. The RT differences between pattern and random high trials gradually increased with practice (shown by the significant ORDER × EPOCH interaction, *F*(4, 204) = 4.46, *p* = .03, _η_^*2*^_*p*_ *=* .08). Other interactions did not reach significance (all *p*s > .37, see Table 3).

**Table 3.** Summary of results from ANOVAs performed on raw RT and standard RT data considering sequence learning.

|  | raw RT |  |  | standard RT |  |  |
| --- | --- | --- | --- | --- | --- | --- |
| | *F* | *p* | $\eta^2_p$ | *F* | *p* | $\eta^2_p$ |
| GROUP | 2.68 | .11 | .05 | 0.08 | .78 | .002 |
| ORDER | 14.48 | < .001** | .22 | 18.79 | < .001** | .27 |
| EPOCH | 35.35 | < .001** | .41 | 4.66 | .001** | .08 |
| GROUP × ORDER | 0.55 | .46 | .01 | 0.001 | .98 | < .001 |
| GROUP × EPOCH | 1.01 | .37 | .02 | 0.88 | .48 | .02 |
| ORDER × EPOCH | 4.46 | .03* | .08 | 0.52 | .72 | .01 |
| GROUP × ORDER × EPOCH | 0.47 | .55 | .01 | 1.03 | .39 | .02 |

Note. \*\* $p < .01$ , \* $p < .05$ , + $p < .10$ .

Although, considering sequence learning, the analysis involving raw RT data did not show differences in average RTs between the groups, for sake of completeness, we conducted identical analyses on standardized RT data as well. Firstly, we compared sequence learning in the last epoch between the groups using mixed-design ANOVA. Similarly to the results on raw RT data, there was no difference in average RTs between the groups (main effect of GROUP, *F*(1, 51) = 0.57, *p* = .46) and participants showed faster RTs on pattern trials compared to random high trials (main effect of ORDER, *F*(1, 51) = 9.71, *p* = .003, _η_^*2*^_*p*_ *=* .16), suggesting significant sequence learning. The analyses did not reveal any group differences in sequence learning (shown by the non-significant ORDER × GROUP interaction, *F*(1, 51) = 0.61, *p* = .44).

To compare sequence learning between the groups on the whole task as well, we used mixed-design ANOVA involving all five epochs of the standard RT data. The results were identical to the ones on raw RT data, overall showing gradual sequence learning (indicated by the significant ORDER × EPOCH interaction, *F*(4, 204) = 4.75, *p* = .023, _η^*2*^ *p*_ *=* .085), but not revealing any group differences (main effect of GROUP and any interaction involving GROUP were not significant, all *p*s > .46, see Table 3).

Performance on *post-block sequence reports*, reflecting explicit sequence knowledge, was also compared between the groups. The independent samples t-test showed that explicit sequence knowledge was greater in the control group than in the stress group (*t*(42.5) = 2.12, *p* = .039, *M*_control_ = 96.2%, *M*_stress_ = 92.4%,).

**Figure 5.**
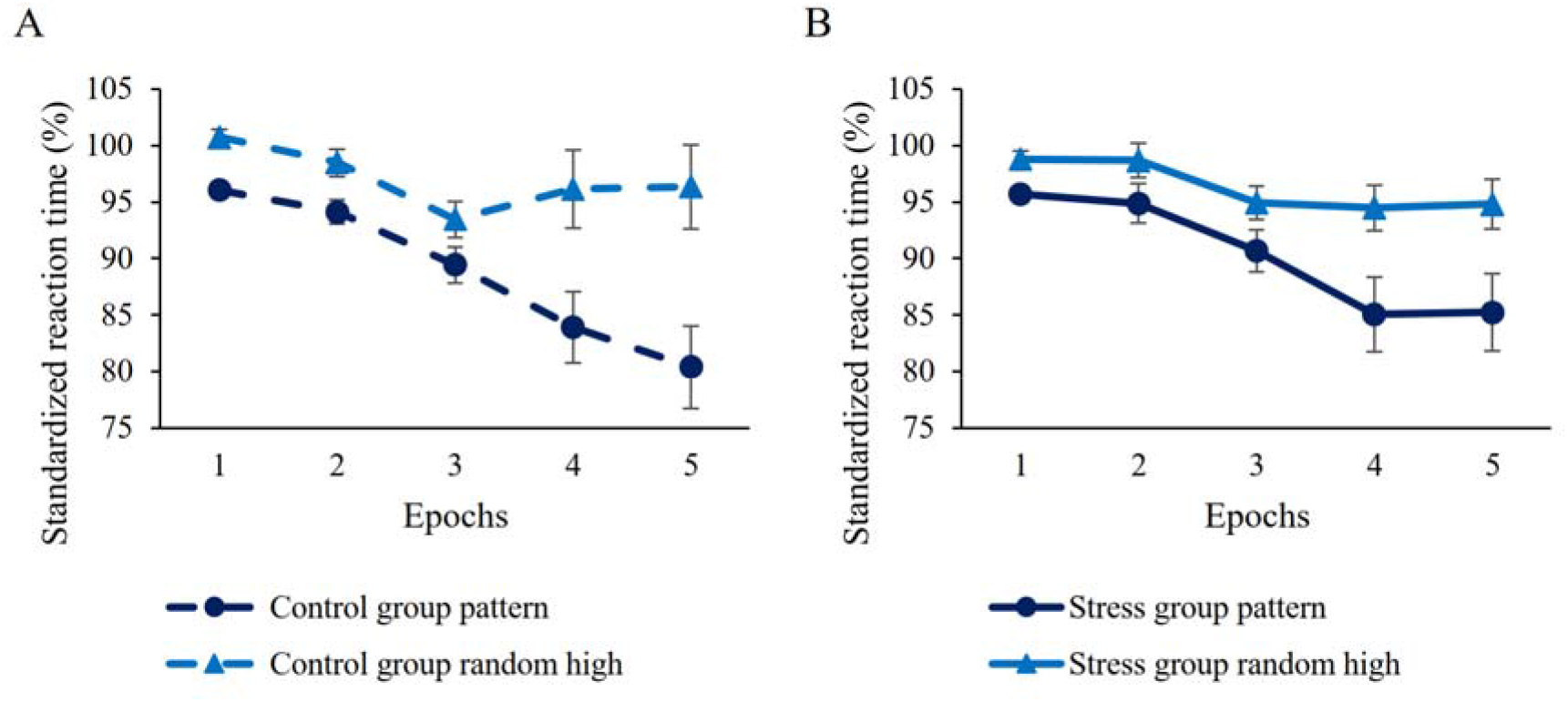
Sequence learning in (A) control and (B) the stress groups. Dashed lines represent the control group, continuous lines represent the stress group. Dark blue lines with circle symbols indicate standardized reaction times on pattern trials, blue lines with triangle symbols indicate standardized reaction times on random high trials. Sequence learning is quantified by the distance between dashed and continuous lines, greater distance represents better learning. Error bars denote the SEM.

## 4. Discussion

To date, only three studies focused on how stress alters the detection and extraction of different types of regularities from the environment (Römer et al., 2011; King, 2013; Dolfen et al., 2019). In the present study, we aimed to get a better understanding of how stress influences the simultaneous acquisition of statistical and sequential regularities. Our results, supported by analyses both in the main text and in the supplementary material, showed that stress affected the acquisition of statistical regularities, while sequence learning was unaffected by the stress induction. Moreover, the stress group showed lower explicit sequence knowledge of the sequence structure presented in the task. Thus, stress promoted an aspect of regularity extraction to some extent while disrupted explicit processes.

Considering statistical learning, to the best of our knowledge, only one study investigated this aspect of regularity extraction under stress. King (2013) found better statistical learning after experimental stress induction on a triplet learning task. In line with this, our study found that stress altered behavioral performance, somewhat boosting the acquisition of statistical regularities. In detail, it seems that stress induction had a more robust effect on the acquisition of statistical regularities at the beginning of the task. There are two possible explanations for this effect, which, at this point, cannot be discriminated. First, acquiring statistical regularities occurs rapidly, often in the early phase of the task, therefore, it is possible that stress boosted the *acquisition* of statistical regularities but not the stabilization of the acquired knowledge. The second explanation is related to the change of cortisol levels during the task. Our analysis showed that cortisol levels were comparable immediately after the ASRT task between the groups. As we did not track cortisol levels during the task, we cannot determine in what pace cortisol levels have changed in the stress group. However, speculatively, cortisol levels might have decreased gradually as the task progressed, leading to differences in learning between the stress and control groups only in the early phase of the task. With these information on hand, we cannot take stand on either explanation, i.e., whether stress only boosts the acquisition but not the stabilization of statistical regularities or stress induction only had a short-term effect, which resulted in group differences only in the early phase of learning. Further studies are needed to directly test this possibility with assessing cortisol levels during the task as well.

As for sequence learning, only two prior studies have investigated the acquisition of sequential regularities under stress. Römer et al. (2011) found delayed sequence learning after pharmacological stress induction. In detail, both the cortisol and placebo groups showed learning, however, the cortisol group showed decreased learning at first, then similar performance to the placebo group at the end of the task. Dolfen et al. (2019) found no effect of stress on deterministic motor sequence learning after experimental stress exposure. Our results fall in line with the latter study’s result as we also have found intact, comparable sequence learning in the stress group after experimental stress induction. As for Römer et al.’s (2011) study, it should be taken into consideration that the authors used pharmacological stress induction (i.e., oral cortisol), therefore, the results may not be clearly transferred into the stress context. There are substantial differences between administrating stress experimentally and pharmacologically, which influences how stress alters learning as well. Pharmacological manipulation often leads to higher glucocorticoid concentration than a stress induction task (Roozendaal et al., 2006; Schwabe et al., 2012). Moreover, stress induction tasks can increase not only the level of cortisol but also trigger the release of many other hormones and neurotransmitters, which can also influence learning (Contarino et al., 1999). Furthermore, pharmacological manipulation does not result in concurrent noradrenergic activity besides glucocorticoid activity, which occurs, however, after inducing stress experimentally (Schwabe et al., 2008). We can speculate that experimental stress induction does not alter deterministic sequence learning while pharmacological induction could decrease it. Since this speculation is based on three studies, further studies are warranted to investigate the effect of stress on the acquisition of sequential regularities in the context of both pharmacological and experimental stress induction.

Note that the cued version of the ASRT task used in this study (Kóbor et al., 2018; Simor et al., 2019) enables us to assess the simultaneous extraction of statistical and sequential regularities within one learning session. The acquisition of statistical regularities occurs incidentally (i.e., without awareness or intention to learn) and relatively rapidly (Kóbor et al., 2018; Simor et al., 2019). In contrast, the acquisition of the sequential regularities shows a gradual trajectory and it can be supported by incidental and intentional learning processes as well, with a typically faster acquisition in the intentional learning condition compared to an incidental learning condition (Howard & Howard, 1997; Howard et al., 2004; Simor et al., 2019). In the cued version of the task, pattern and random elements are visually distinguishable, and participants are instructed to find the sequence, which creates a partly intentional learning condition. However, it is important to note that even in an intentional learning situation, RTs tend to reflect a more implicit, incidental measure of sequence knowledge, while explicit sequence knowledge measured by post-block sequence reports tends to reflect a more intentional process. As typical response times are under 500 ms, the fast pace of the task makes it unlikely that explicit sequence knowledge would have an impact on participants’ RTs. Therefore, the consciously accessible knowledge of the sequence (reflected by the post-block sequence report score) and sequence learning (reflected by the RTs) seems to be dissociable measures (Horváth et al., 2019). The sequence report may be considered as a more explicit measure of sequence knowledge, while the sequence learning RT scores may serve as an implicit measure of sequence acquisition, even in an intentional learning condition. In this view, in the present study, stress disrupted the explicit acquisition of the sequence knowledge, while did not alter the implicit measure of sequence learning.

In sum, the present study investigated whether the simultaneous acquisition of two types of regularities, namely statistical and sequential regularities, is influenced by experimental stress induction. Our results showed that stress affected the acquisition of statistical regularities as participants in the stress group showed higher statistical learning compared to the control group. Stress did not alter sequence learning but disrupted the explicit acquisition of the repeating sequence. Our results fall in line with previous studies examining the effect of stress on either statistical or sequence learning (King, 2013; Dolfen et al., 2019), however, we went beyond these studies by investigating these learning processes in one experimental design. Moreover, we can interpret our results from an evolutionary point of view: in a risky, dangerous situation, the human brain extracts statistical regularities embedded in our environment faster. This stress-related boost provides an advantage for the individual to effectively manage the situation.

## Acknowledgements

This research was supported by the National Brain Research Program (project 2017-1.2.1-NKP-2017-00002); Hungarian Scientific Research Fund (NKFIH-OTKA K 128016, to D.N., NKFIH-OTKA PD 124148 to K.J.); János Bolyai Research Scholarship of the Hungarian Academy of Sciences (to K.J.); IDEXLYON Fellowship of the University of Lyon as part of the Programme Investissements d’Avenir (ANR-16-IDEX-0005) (to D.N.). We thank Kate Schipper for her comments and suggestions on the previous version of the manuscript.

## Supplementary materials

### 1. Supplementary data analyses on sample with more lenient exclusion criteria

As the number of participants not following task instructions is 10% of all participants, we decided to show the analyses including these seven participants in the Supplementary Material. The analyses are identical to the ones shown in the manuscript. The sample consisted of 60 participants (16 men, 44 women) with a mean age of 20.88 (*SD* = 1.68). The stress group consisted of 31 participants (8 men, 23 women) and the control group consisted of 29 participants (8 men, 21 women). 13 women in the stress group and 8 women in the control group took oral contraceptives regularly.

#### 1.1. Supplementary Results

##### 1.1.1. The effectiveness of stress induction

We used both objective (cortisol levels) and subjective (questionnaire-based ratings) measurements to test the effectiveness of stress induction. For the *objective measurement*, the ANOVA confirmed the effectiveness of the stress induction. We found a significant TIME × STRESS EXPOSURE interaction (*F*(2, 116) = 28.06, *p* < .001, _η_^*2*^_*p*_ *=* .326). The post hoc analysis revealed that cortisol levels in the stress and control groups differed 15 minutes after the stress/control procedure (t2; *M*_stress_ = 32.90, *SD*_stress_ = 15.90, *M*_control_ =21.89, *SD*_control_ = 19.17, *p* = .018), whereas immediately before the stress/control procedure (t1) and immediately after the ASRT task (t3) the two groups’ cortisol levels did not differ significantly (both *p*s > .42, Supplementary Figure 1). For the *subjective ratings* of affective state after stress induction or control task, the t-tests showed significantly higher subjective stress, pain and unpleasantness levels in the stress group compared to the control group (see Supplementary Table 1).

**Supplementary Table 1.** Descriptive data of the participants’ subjective stress measurements.

|  | Group |  |  |  | *t* | *p* |
| --- | --- | --- | --- | --- | --- | --- |
|  | Stress |  | Control |  |  |  |
|  | *(n = 31)* |  | *(n = 29)* |  |  |  |
|  | *M* | *SD* | *M* | *SD* |  |  |
| Subjective ratings of stress | 31.19 | 26.28 | 8.51 | 11.98 | -4.24 | <.001 |
| Subjective ratings of unpleasantness | 59.29 | 29.17 | 12.20 | 14.02 | -7.87 | <.001 |
| Subjective ratings of painfulness | 60.00 | 24.89 | 1.20 | 5.61 | -12.42 | <.001 |

*Note.* Immediately after the stress induction or control task, participants rated how stressful, unpleasant and painful (where 0 = not at all, 100 = very) the stress induction or control task was.

**Supplementary Figure 1.**
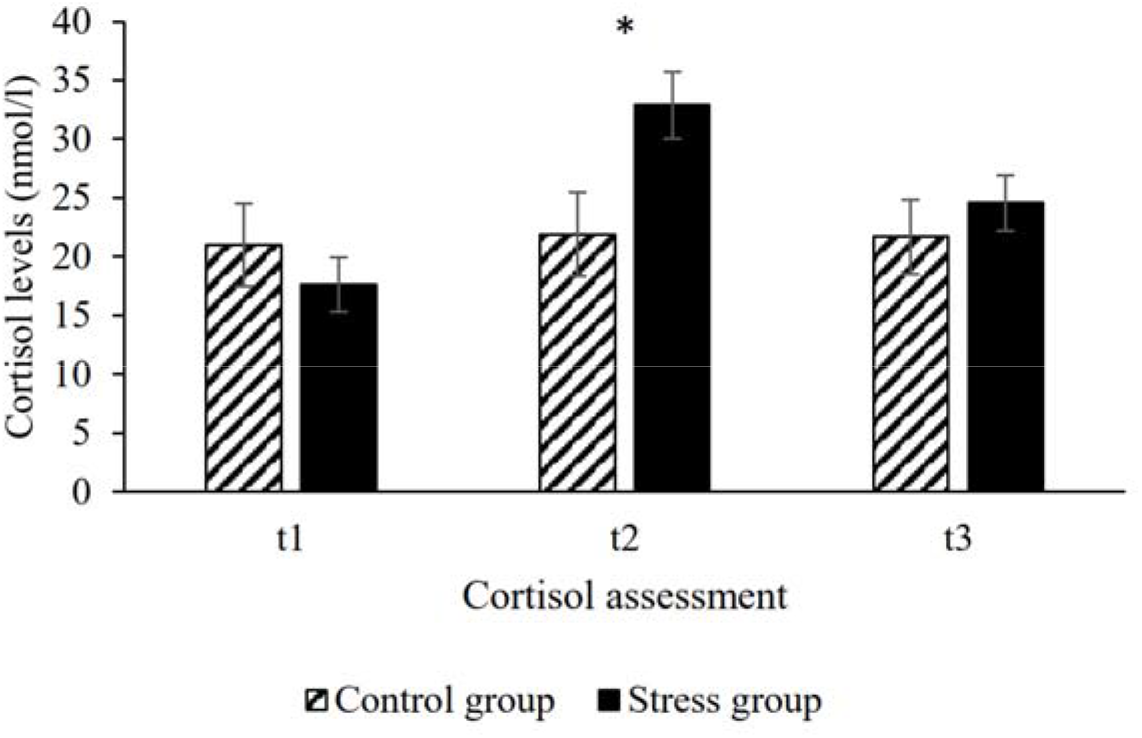
The effectiveness of stress induction. Salivary cortisol levels were assessed immediately before (t1), 15 minutes after (t2) and 45 minutes after (t3) the stress induction or control task. Error bars denote standard error of mean.

##### 1.1.2. General motor performance

To assess general motor execution, a mixed design ANOVA was conducted (see Data analysis in the manuscript). The ANOVA revealed that participants became faster as the task progressed (main effect of BLOCK, *F*(2, 116) = 67.09, *p* < .001, _η_^*2*^_*p*_ *=* .54). Importantly, no group differences was found (main effect of GROUP, *F*(1, 58) = 0.01, *p* = .93; GROUP × BLOCK interaction, *F*(2, 116) = 0.10, *p* = .88).

##### 1.1.3. The effect of stress induction on statistical learning

As statistical learning is a rapid process (Kóbor et al., 2018; Simor et al., 2019), we focused on the beginning of the task. At first, we examined statistical learning only in the first epoch using a mixed-design ANOVA (see section 2.5 in the manuscript) between the stress and control groups. Participants had faster responses to random high than random low trials (shown by the significant main effect of PROBABILITY, *F*(1, 58) = 31.19, *p* < .001, _η_^*2*^_*p*_ *=* .35). Over trials, there was no difference in reaction times between the groups (main effect of GROUP, *F*(1, 58) = 1.13, *p* = .29). Importantly, the stress group showed higher statistical learning (i.e., greater RT difference between random high and random low trials) than the control group (indicated by the significant PROBABILITY × GROUP interaction, *F*(1, 51) = 5.53, *p* = .02, _η_^*2*^_*p*_ *=* .09, stress group: *M*_*random high*_ = 378.48 ms, *M*_*random low*_ = 396.55 ms, statistical learning score: *M* = 18.07 ms; control group: *M*_*random high*_ = 398.28 ms, *M*_*random low*_ = 405.64 ms, statistical learning score: *M* = 7.36 ms).

To investigate statistical learning in the stress and control groups on the whole task, we conducted a mixed-design ANOVA on the RTs (see section 2.5 in the manuscript). Over groups, RTs were faster on random high trials than on random low trials (shown by the significant main effect of PROBABILITY, *F*(1, 58) = 155.70, *p* < .001, _η_^*2*^_*p*_ *=* .73), indicating significant statistical learning. Participants became faster with practice on both trials (inidcated by the significant main effect of EPOCH, *F*(4, 232) = 6.07, *p* = .01, _η_^*2*^ *p =* .095).

There were no differences in average RTs between the groups (main effect of GROUP, *F*(1, 58) = 1.74, *p* = .19). The RT differences between random high and random low trials increased with practice (indicated by the significant PROBABILITY × EPOCH interaction, *F*(4, 232) = 2.85, *p* = .03, _η_^*2*^ *p =* .05). Importantly, the time course of statistical learning was different between the groups (indicated by the marginally significant GROUP × PROBABILITY × EPOCH interaction, *F*(4, 232) = 2.08, *p* = .08, _η_^*2*^_*p*_ *=* .04, see Supplementary Figure 2). Follow-up analysis on the statistical learning score revealed a significant difference in the first epoch: the stress group showed higher statistical learning than the control group (stress: *M* = 18.07 ms, control: *M* = 7.36 ms, *p* = .02). Additionally, the stress group showed marginally higher statistical learning in the third epoch as well (stress: *M* = 24.11 ms, control: *M* = 15.03 ms, *p* = .08), while there were no group differences in the remaining epochs (all *p*s > .22). Other interactions did not reach significance (all *p*s > .17).

**Supplementary Figure 2.**
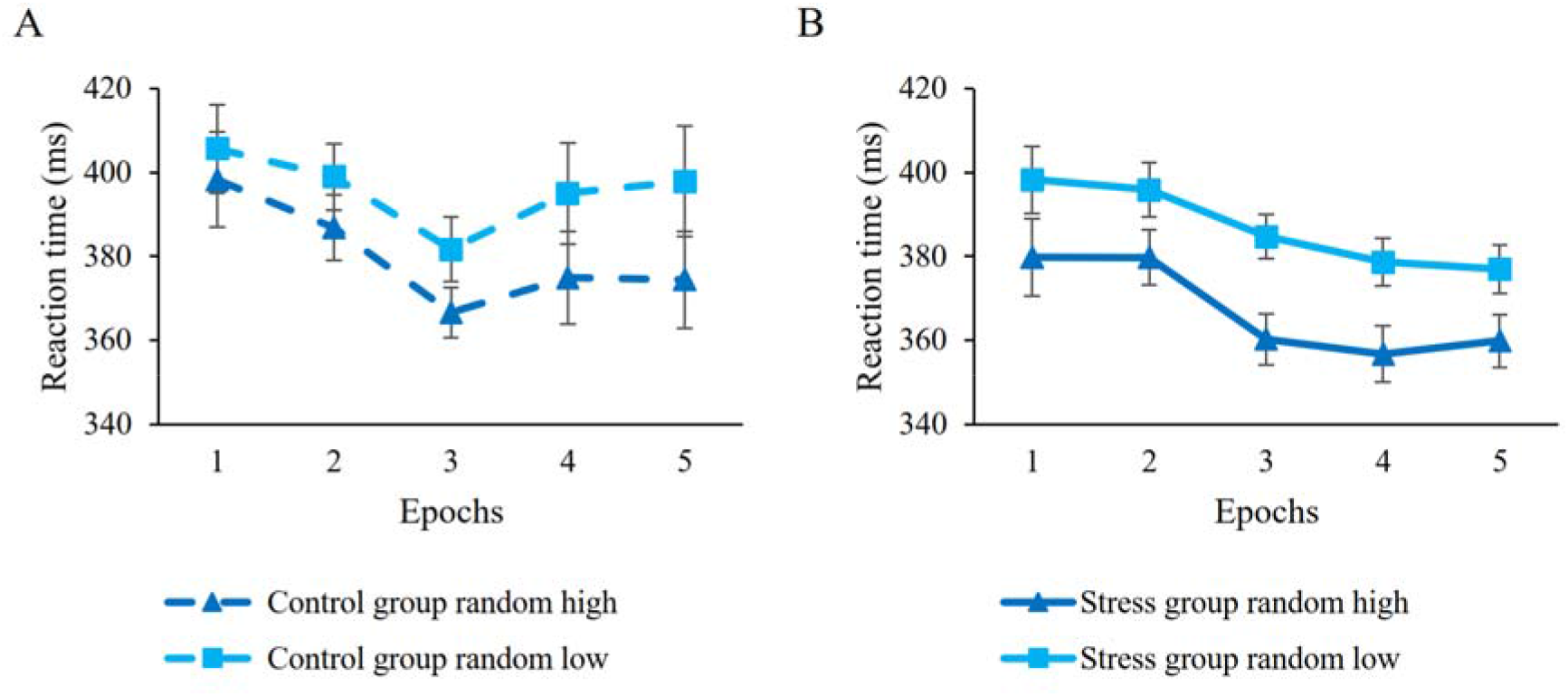
Statistical learning in (A) control and (B) the stress groups. Dashed lines represent the control group, continuous lines represent the stress group. Darker blue lines with triangle symbols indicate reaction times on the random high trials, light blue lines with square symbols indicate reaction times on the random low trials. Statistical learning is quantified by the distance between dashed and continuous lines, greater distance represents better learning. Error bars denote standard error of mean.

##### 1.1.4. The effect of stress induction on sequence learning

As sequence learning is a gradual process (Kóbor et al., 2018; Simor et al., 2019), at first, we investigated sequence learning between the groups in the last epoch using a mixed-design ANOVA (see section 2.5 in the manuscript). Participants showed faster responses to pattern than to random high trials (main effect of ORDER, *F*(1, 58) = 9.41, *p* = .003, _η^*2*^ *p*_ *=* .14). Over trials, participants showed similar reaction times (main effect of GROUP, *F*(1, 58) = 0.50, *p* = .48). The analysis showed comparable sequence learning between the groups (shown by the non-significant ORDER × GROUP interaction, *F*(1, 58) = 0.58, *p* = .45).

To examine sequence learning on the whole task, we also used mixed-design ANOVA on the RT (see section 2.5 in the manuscript). Participants showed faster RTs on pattern trials compared to random high trials (indicated by the main effect of ORDER, *F*(1, 58) = 14.95, *p* < .001, _η_^*2*^ *p =* .21), indicating significant sequence learning. With practice, participants showed faster RTs on both trials (shown by the significant main effect of EPOCH, *F*(4, 232) = 41.61, *p* < .001, _η_^*2*^_*p*_ *=* .42). Average RTs did not differ between the groups (main effect of GROUP, *F*(1, 58) = 1.24, *p* =.27). Moreover, the RT differences between pattern and random high trials gradually increased with practice (shown by the significant ORDER × EPOCH interaction, *F*(4, 232) = 4.05, *p* = .04, _η_^*2*^ *p =* .07, see Supplementary Figure 3). Other interactions did not reach significance (all *p*s > .47).

Performance on *post-block sequence reports*, reflecting explicit sequence knowledge, was also compared between the groups. The independent samples t-test showed similar explicit sequence knowledge in the groups (*t*(58) = 0.65, *p* = .514, *M*_control_ = 89.5%, *SD*_control_ = 21.0%, *M*_stress_ = 86.1%, *SD*_stress_ = 18.1%).

**Supplementary Figure 3.**
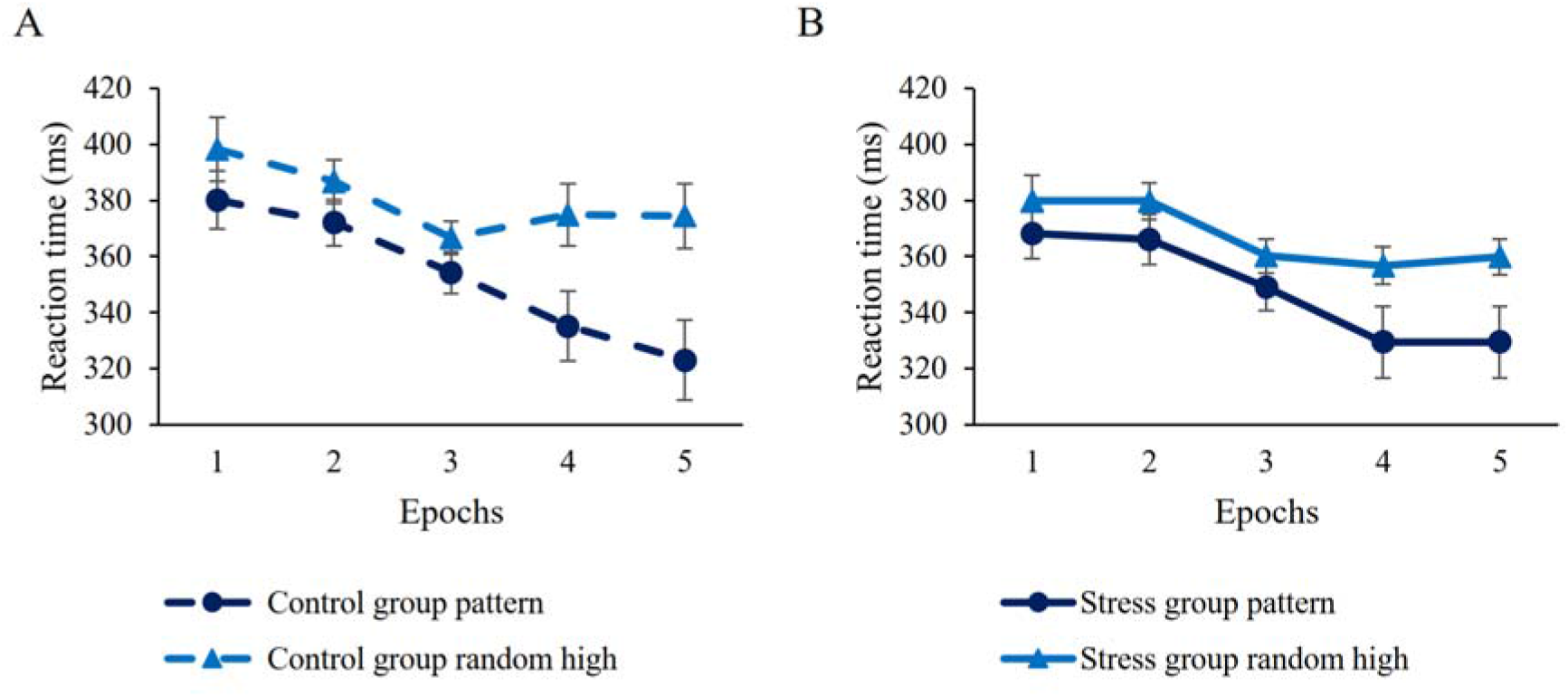
Sequence learning in (A) control and (B) the stress groups. Dashed lines represent the control group, continuous lines represent the stress group. Dark blue lines with circle symbols indicate reaction times on pattern trials, lighter blue lines with triangle symbols indicate reaction times on random high trials. Sequence learning is quantified by the distance between dashed and continuous lines, greater distance represents better learning. Error bars denote standard error of mean.

